# MASST: A Web-based Basic Mass Spectrometry Search Tool for Molecules to Search Public Data

**DOI:** 10.1101/591016

**Authors:** Mingxun Wang, Alan K. Jarmusch, Fernando Vargas, Alexander A. Aksenov, Julia M. Gauglitz, Kelly Weldon, Daniel Petras, Ricardo da Silva, Robby Quinn, Alexey V. Melnik, Justin J.J. van der Hooft, Andrés Mauricio Caraballo Rodríguez, Louis Felix Nothias, Christine M. Aceves, Morgan Panitchpakdi, Elizabeth Brown, Francesca Di Ottavio, Nicole Sikora, Emmanuel O. Elijah, Lara Labarta-Bajo, Emily C. Gentry, Shabnam Shalapour, Kathleen E. Kyle, Sara P. Puckett, Jeramie D. Watrous, Carolina S. Carpenter, Amina Bouslimani, Madeleine Ernst, Austin D. Swafford, Elina I. Zúñiga, Marcy J. Balunas, Jonathan L. Klassen, Rohit Loomba, Rob Knight, Nuno Bandeira, Pieter C. Dorrestein

## Abstract

We introduce a web-enabled small-molecule mass spectrometry (MS) search engine. To date, no tool can query all the public small-molecule tandem MS data in metabolomics repositories, greatly limiting the utility of these resources in clinical, environmental and natural product applications. Therefore, we introduce a **Ma**ss **S**pectrometry **S**earch **T**ool (MASST) (https://proteosafe-extensions.ucsd.edu/masst/), that enables the discovery of molecular relationships among accessible public metabolomics and natural product tandem mass spectrometry data (MS/MS).

---

The ability to discover related sequences of proteins or genes in publicly accessible sequence data using Basic Local Alignment Search Tool (BLAST), connected to public sequence data repositories through a web interface (WebBLAST, https://blast.ncbi.nlm.nih.gov/Blast.cgi), was introduced in the 1990s.^1^ It has garnered more than 138,159 citations according to Google Scholar, placing it among the most widely used bioinformatics tools. WebBLAST enabled detection of the number of sequences in public repositories related to a given query, the organisms in which those sequences occur, and the evolutionary and inferred functional relationships among related sequences. It therefore permitted a broad community to answer simple but scientifically compelling questions such as: Is a protein or DNA sequence common or rare? How is this sequence distributed among different kinds of organisms? What other sequences are related to this sequence (evolutionary variants, or new mutations, or synthetic constructs)? In the early days of making DNA or protein sequence data publicly available, the “metadata” (e.g., contextual information about the sample, population and location the sequence came from, and technical information about how it was produced) in the public repositories was limited and no standards existed. This is a situation similar to the current status of much of the mass spectrometry data in the public domain. However, when publicly deposited data has metadata available, such as organism, location of sampling, host phenotypes such as diseases, etc., it becomes possible to start building higher-level hypotheses regarding the evolutionary, ecological or functional relationships among these DNA, RNA or protein sequences. The development of the ability to search data with added context continues to have profound impacts on fields including medicine, chemistry, genetics, molecular biology, genomics, microbiology, and ecology.

Algorithms developed for mass spectrometry data, including molecular networking^2^ and fragmentation trees^3^, enable similarity searches, while powerful metabolomics analysis software infrastructures, such as MS-DIAL^4^, MetaboAnalyst^5^, XCMS Online^6^, HMDB^7^, some of which have been available for over a decade, focus on annotation of MS/MS spectra or finding statistical relationships between molecular features. However, none of the existing tools enable searching against public data in repositories. Finding the distribution of specific data of interest, in this case MS/MS spectra, including unannotated spectra and structural analogs among data in public metabolomics’ and natural product’s data mass spectrometry data repositories, is not yet possible. Consequently, there is limited appreciation for the potential of mass spectrometry to benefit from the transformative features equivalent to those described above for sequence data. Deposition of untargeted mass spectrometry data in the public domain is experiencing rapid growth, from 910 metabolomics datasets available on March 2017^8^ to more than 2,000 downloadable metabolomics datasets in January 2019 (about half of which have MS/MS data).^9^ Despite the increasingly growing accessibility of metabolomics and natural products data, including environmental and clinical mass spectrometry datasets, public small molecule mass spectrometry data itself has seen little reuse.^10^ Therefore, we introduce MASST to enable the reuse of publicly available untargeted tandem mass spectrometry data, akin to the way WebBLAST interfaced with public sequence repositories enables online searching of public sequencing data.

To provide metabolomics MS/MS search capabilities similar to those that have been available to the sequencing community for almost 30 years, we engineered a web-based system that enables the searching of data deposited in the public data repository portion of the GNPS/MassIVE knowledge base^11^ and an analysis infrastructure for a single MS/MS spectrum. The developments required for enabling MASST searches included converting public data to a uniform open format^12^ (irrespective of instrument type and original data format), the ability to trace the original file where each MS/MS spectrum originated, and a reporting infrastructure that provides all identical or similar MS/MS spectra found in the public data along with their associated metadata. Key reasons why MASST has now become possible, and not ∼30 years ago when the sequencing community launched WebBLAST, are: 1) only in recent years have a sufficient amount of small molecule untargeted mass spectrometry datasets become publicly available to warrant the development of such capabilities (∼1,100 untargeted datasets and ∼110,000,000 spectra in ∼150,000 files as of Dec 11, 2018), 2) increased adoption of universal, non-vendor specific MS data formats makes this the first time that many publicly available datasets have been converted to the same data format ^28, 29^ and 3) the engineering of the infrastructure to enable tracking of information of all public data (in GNPS/MassIVE) and connecting each MS/MS spectrum to its own unique metadata entries had not been developed yet.

Entering data from a single MS/MS spectrum would be the equivalent to entering a single protein or gene sequence into WebBLAST. Akin to WebBLAST, with MASST, it is possible to search against multiple repositories, including GNPS/MassIVE ^11^, Metabolomics Workbench ^13^, Metabolights ^14^ or the non-redundant (nr) MS/MS library of all unique MS/MS spectra from all three repositories combined. MASST searching using multiple repositories was made possible by converting data uploaded to the Metabolomics Workbench and MetaboLights repositories to the same open mass spectrometry format within the GNPS/MassIVE data storage environment. MASST searches and the data retrieval infrastructure report results within a user defined similarity score to the input data. The report returns the origin of the matched MS/MS spectrum with respect to the dataset and file information, and any sample information orother metadata associated with the file (when available) (**Figure 1 a-e**). Further, datasets and files can be tagged with sample or spectral information by the community of MASST users that then becomes a part of the metadata reported back in a MASST search. Approximately 23,000 additional files with ∼230,000 tags, mostly human-associated, have been manually curated by the authors of this paper, thus providing a foundation for reporting MASST searches.

**Figure 1.**
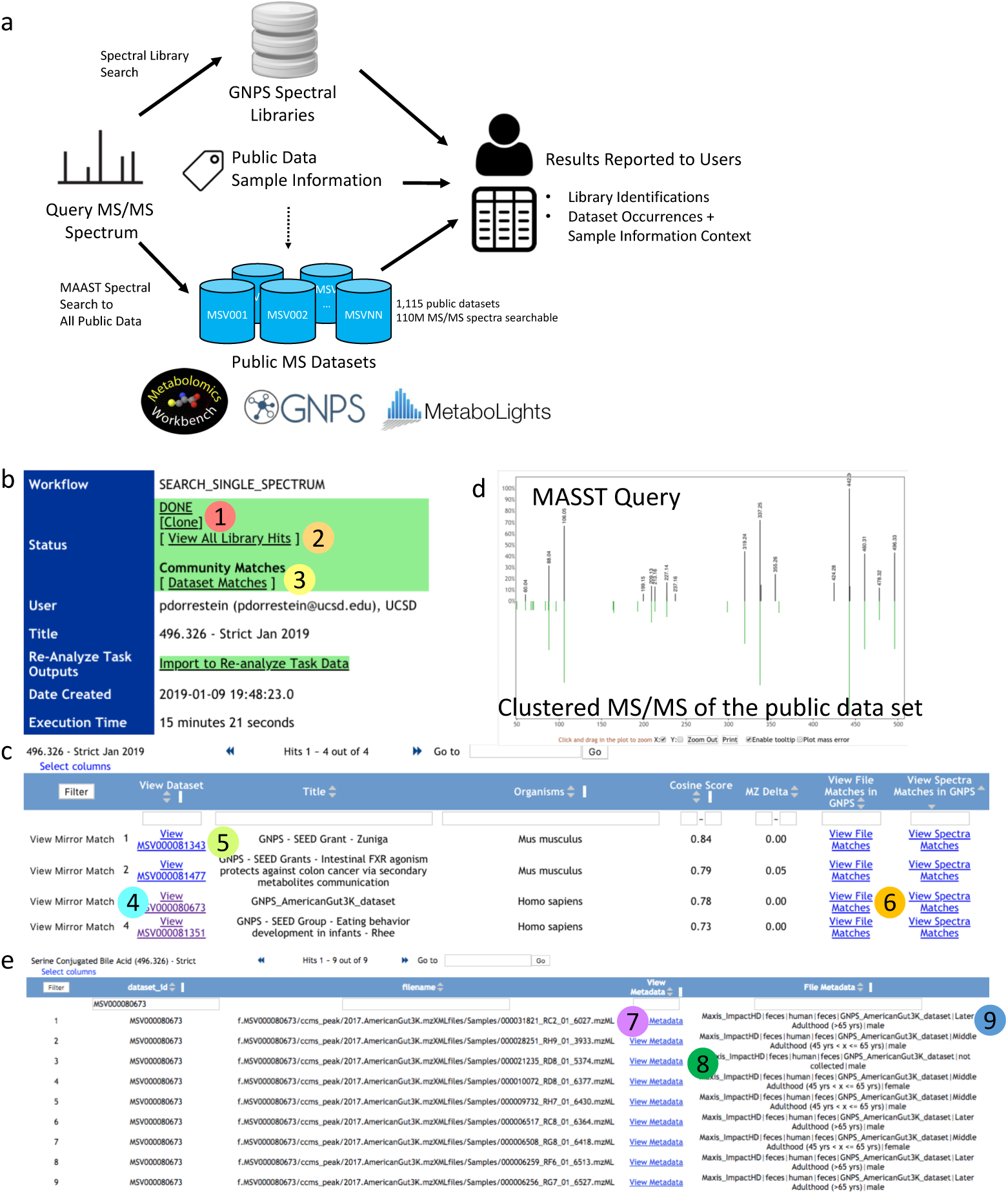
MASST search, reporting and inspection of matches. **a.** Overview of MASST query procedure. MASST queries MS/MS spectra across all public metabolomics data, including those deposited at GNPS, Metabolomics Workbench, and Metabolights. Combining these matches with sample information at the file granularity and spectral library search, provides users with a report about MS2 compound annotation and MS2 sample information context. Once a MASST search is completed via https://proteosafe-extensions.ucsd.edu/masst/, the results can be found under the users job tab or via the link provided in an email. **b.** shows the opening page. There are two options (**2** and **3**) for inspecting the data and options for cloning a job (**1**). Clicking (**2**) will reveal all MS/MS spectral matches within the user defined settings. There can be none, one or more than one match for a given input spectrum. **c.** Clicking (**3**) will reveal all data sets that contain an MS/MS spectrum that has a match to the input spectrum and sample information associated with that data set. **d**. clicking on “View Mirror Match” (**4**) shows the mirror match between the input spectrum and the merged MS/MS spectrum enabling manual inspection of this match; “View MSV0000…..” (**5**) will bring you to the data set and all uploaded information associated with this data set can be found or is linked in this location. (**6**) opens up the file information window and tabulated metadata. (**7**) shows the files where MS/MS matches are found while (**8**) Link-out to full sample information for the file. (**9**) are the abbreviated (and filterable) sample information associated with the files. If no sample information has been uploaded with the original data, then this will be blank. The MASST_GNPS job link for this search to enable the reader to navigate the same results can be found here. https://gnps.ucsd.edu/ProteoSAFe/status.jsp?task=bac3d3788e704af59e4a15a5146e4d6b

The report returned by MASST also includes matches to any reference spectra in public MS/MS spectral libraries if the matches are within the search parameters specified by the user. Searched libraries include GNPS user contributed spectra^11^, GNPS libraries^11^, all three MassBanks^15,16,17^, ReSpect^18^, MIADB/Beniddir^27^, Sumner/Bruker, CASMI^20^, PNNL lipids^21^, Sirenas/Gates, EMBL MCF and several other libraries that can be found here: https://gnps.ucsd.edu/ProteoSAFe/libraries.jsp. The report further enables interactive inspection of the results and displays the matches as a mirror view (**Figure 1c**) akin to the way that NCBI’s BLAST web server provides a picture of the aligned sequences to guide the exploration of the such alignment results. The GNPS/MassIVE upload portal accepts metadata information at the dataset level, file level and single annotated spectrum level. Examples of sample information include (i) instrument type, phylogeny (according to NCBI taxonomy) and keywords at the dataset level, (ii) phylogeny, sample type, age, sex, body site (defined using the Uberon anatomy ontology^22^), and disease^23^ at the file level, and (iii) source, biological activity, and structural class information at the single annotated spectrum level. In addition, GNPS/MassIVE is compatible with metadata formats from other software tools, e.g. QIIME2 and Qiita, which are used to analyze microbiome data and have a large controlled vocabulary that can be imported.^24,25,26^ Further, the sample information uploaded to the other repositories is also made accessible *via* the MASST report.

As is the case with all early and current sequence repositories, there is still limited information at the dataset and file level, but the metadata that is already present in the public domain can provide insight into the specific MS/MS signals being investigated (see Box 1 for representative examples of usage). Although the amount and quality of metadata is growing, datasets do not always have sufficiently detailed metadata. This is why GNPS enables re-annotation of multiple levels of metadata as the community knowledge increases, while retaining provenance of all changes.^11^ When insufficient metadata is available for interpretation of the public dataset search results, then the original depositors of the public data can be contacted, something that is still often necessary within the sequencing community. This is also expected with MASST and could foster collaborations worldwide.

Similar to the NCBI’s WebBLAST server, the use of MASST is designed to be straightforward. MASST can be accessed in three ways: 1) direct access via https://proteosafe-extensions.ucsd.edu/masst/, by copying/pasting the MS/MS spectrum peak list reported as *m/z* and intensity separated by a space for each fragment ion (aka product ion), 2) input of files, in open mass spectrometry formats, such as .mzML, .mzXML, .MGF, or 3) automated entry of MS/MS of interest from online GNPS data analysis. Manual entry provides the researcher the ability to enter data from theoretical spectra or other spectra found in published papers or supporting information without the original experimental data. Option (3) allows users to launch a MASST search via direct links provided in the molecular network version 2.0 output created within the GNPS infrastructure^11^, which automatically redirects to the MASST search page with the prepopulated spectral data. When performing feature-based molecular networking, this option can be found under the “View all cluster ID” (shows all spectrum clusters in the search, regardless o whether they’re identified) and then clicking “Search spec”. The MS/MS spectrum provided via the MASST website or as a link-out from a GNPS search is then searched against all public data with user defined parameters of minimum number of ions to match, precursor (parent) and product (fragment) ion tolerances, and analog similarity searches based on non-identical precursor masses.^2^ MASST searches retrieve all associated sample information (dataset and files) that match the MS/MS input spectrum query. A typical search takes about 10-20 min, with multiple searches queries are placed in a queue for parallel execution as resources become available. To promote data analysis reproducibility, the results of the job are stored in each user’s space and can be found under the “Jobs” tab accessible through the banner in the GNPS browser (http://gnps.ucsd.edu). Only MASST jobs run while logged in will be retained. The search parameters are also retained with each job and constitute a provenance record that can be provided as hyperlinks to share with others, e.g. collaborators and in publications. These jobs can be shared, cloned, and rerun with or without alterations of the input parameters (examples of links to jobs provided in Box1). This could lead to new additional matches in case relevant public data was uploaded since the last time a MASST search was done. The matches of MS/MS spectra among datasets are the equivalent to level two (putative annotation based on spectral library similarity) or three (putatively characterized compound class based on spectral similarity to known compounds of a chemical class) according to the 2007 metabolomics standards initiative^26^. Similar to short sequence reads, MASST searches will currently not distinguish chemicals that have nearly identical fragmentation patterns, such as isomeric compounds, which would require an authentic standard and the use of an orthogonal property (such as the retention time).

In some cases when a MASST search returns no matches, it may be because there is no data that matches or it is possible that MS/MS matches in the currently available public datasets fall outside the specified search parameters. MASST should be used with these caveats in mind when formulating a hypothesis. It is expected that MASST can be used for a wide array of applications, analogous to BLAST. Uses of MASST are expected to range from translation of experiments *in vitro* or in model organisms to humans, to asking broad ecological questions. In Box 1, we have provided ten examples that highlight the types of discoveries that users may make only by searching across all public data and we expect that the user community will come up with additional innovative ways to use MASST.

## Contributions

PD and MW came up with the concept of MASST. MW and NB performed the engineering to enable MASST. MW, AKJ, JVDH, JMG, MP, EOE, KW, CMA, FDO, EB, AB curated metadata. FV, JMG, LLB, KW, EB, AA and CSC generated data for the manuscript. EG synthesized the bile acids. PD, MW, DP, JDW, MJ, LFN, JMG, EIZ, LLB, KEK, SPP, AMCR, FV, KW, AA performed experiments and/or analysis for Box 1. PD, DP, LFN, JVDH, JMG, AA, AMCR, FV, KW, AB, FDO, ME, RS tested the MASST infrastructure and downloaded public data. PD, NB, EIZ, RL, RK, ADS, MJB, JLK provided supervision and funding for the project. PD, AJ, DP, JVDH, ME, JMG, AA, AMCR, RK, JLK, LF, NB, MW wrote and edited the manuscript.

## Acknowledgements

The conversion of the data from different repositories was supported by R03 CA211211 on reuse of metabolomics data, the development of a user-friendly interface was in part supported by Gordon and Betty Moore Foundation through Grant GBMF7622. The UC San Diego Center for Microbiome Innovation supported the campus wide SEED grant awards for data collection that enabled the development much of this infrastructure. AKJ thanks the American Society for Mass Spectrometry for the 2018 Postdoctoral Career Development Award. We further acknowledge Claire O’Donovan and Kenneth Haug for help with navigating the MetaboLights data repository. JVDH was supported by a ASDI eScience grant (ASDI.2017.030) from the Netherlands eScience Center (NLeSC). EIZ and LLB were supported by NIH grants AI081923 and AI113923. AMCR, KEK, SPP, JLK, MJB, and PCD were supported by NSF grant IOS-1656475. AB was supported by National Institute of Justice Award 2015-DN-BX-K047. FV was supported by the Department of Navy, Office of Naval Research Multidisciplinary University Research Initiative (MURI) Award, Award number N00014-15-1-2809. DP was supported by the German Research Foundation (DFG) with Grant PE 2600/1. Additional support for data acquisition and data storage was provided by P41 GM103484 Center for Computational Mass Spectrometry, Instrument support though NIH S10RR029121, RL is supported by NIH grants R01DK106419, 5P42ES010337, and 5UL1TR001442, NIH K01DK116917 to J.D.W. The development of the web interface and harmonization with Qiita was in part supported by the Sloan Foundation.

---

Box 1 (could become SI). Below are ten representative examples to illustrate how the community can use MASST to address scientific questions

**1) Are specific molecular features detected via mass spectrometry in one clinical cohort also observed elsewhere?** Nonalcoholic fatty liver disease (NAFLD) is the leading cause of chronic liver disease in the United States, afflicting 80-100 million Americans. It can be broadly sub-divided into two categories: Nonalcoholic fatty liver (NAFL), which is the non-progressive form of NAFLD with minimal risk of progression to cirrhosis, and nonalcoholic steatohepatitis (NASH), with substantial risk of progression to cirrhosis.^1^ The fecal samples of individuals with NAFLD with and without advanced fibrosis (n=28) with corresponding healthy controls of close relatives (twins, siblings-siblings and parents-offspring, n=110) were analyzed using untargeted data dependent mass spectrometry. Partial least squares-discriminant analysis, Random Forest and other statistical methods consistently suggested that a mass spectrometry feature with *m/z* 540.3677 was most strongly associated with diagnosed NAFLD (Box Figure 1a). We had no other knowledge about this molecular feature other than its precursor mass and mass fragments. MASST revealed that the MS/MS spectrum associated with this feature was found in 27 datasets (9 mouse fecal, 2 rat fecal, 1 *Escherichia coli* and 15 human fecal studies). Deeper inspection of the datasets revealed 2 studies of mouse models of NAFLD. MASST allows one to find these datasets that can be investigated further. Upon re-exploring the data for one of the latter studies^1^, this molecule was found in higher abundance in mice fed a high-fat diet, a condition that induced NAFLD in these animals thus supporting the discovery that this molecule may indeed be NAFLD-related (Box Figure 1 b and c). Furthermore, this molecule disappeared when animals were treated with antibiotics^1^, suggesting a microbial origin of this molecule. To gain additional insight, the data were subjected to molecular networking which revealed that the MS/MS spectrum was related chenodeoxycholate but with a mass shift of phenylalanine. This putative molecule would be similar to the related to microbially derived cholyphenylalanine by a loss of an oxygen.^2^ As the mass spectrometry is largely blind to regiochemistry, four different phenylalanine amidate conjugates were synthesized. The phenylalanine conjugate of the deoxycholic acid co-migrated with this feature, classifying the annotation as level one according to the 2007 Metabolomics Standards Initiative (Box Figure 1e).^3^

Taken together, these findings constitute a discovery that the most significant differentiating molecule associated with NAFLD in fecal samples is a microbially-derived bile acid. This molecule is observed in the human cohort, but is also recapitulated in the murine model of the disease. This serves as an important confirmation that did not require any additional experimental analysis due to availability and reuse of existing public data.

MASST_GNPS job link: https://gnps.ucsd.edu/ProteoSAFe/status.jsp?task=9a703332aa794074bbb58a148d32cfff

**Box Figure 1.**
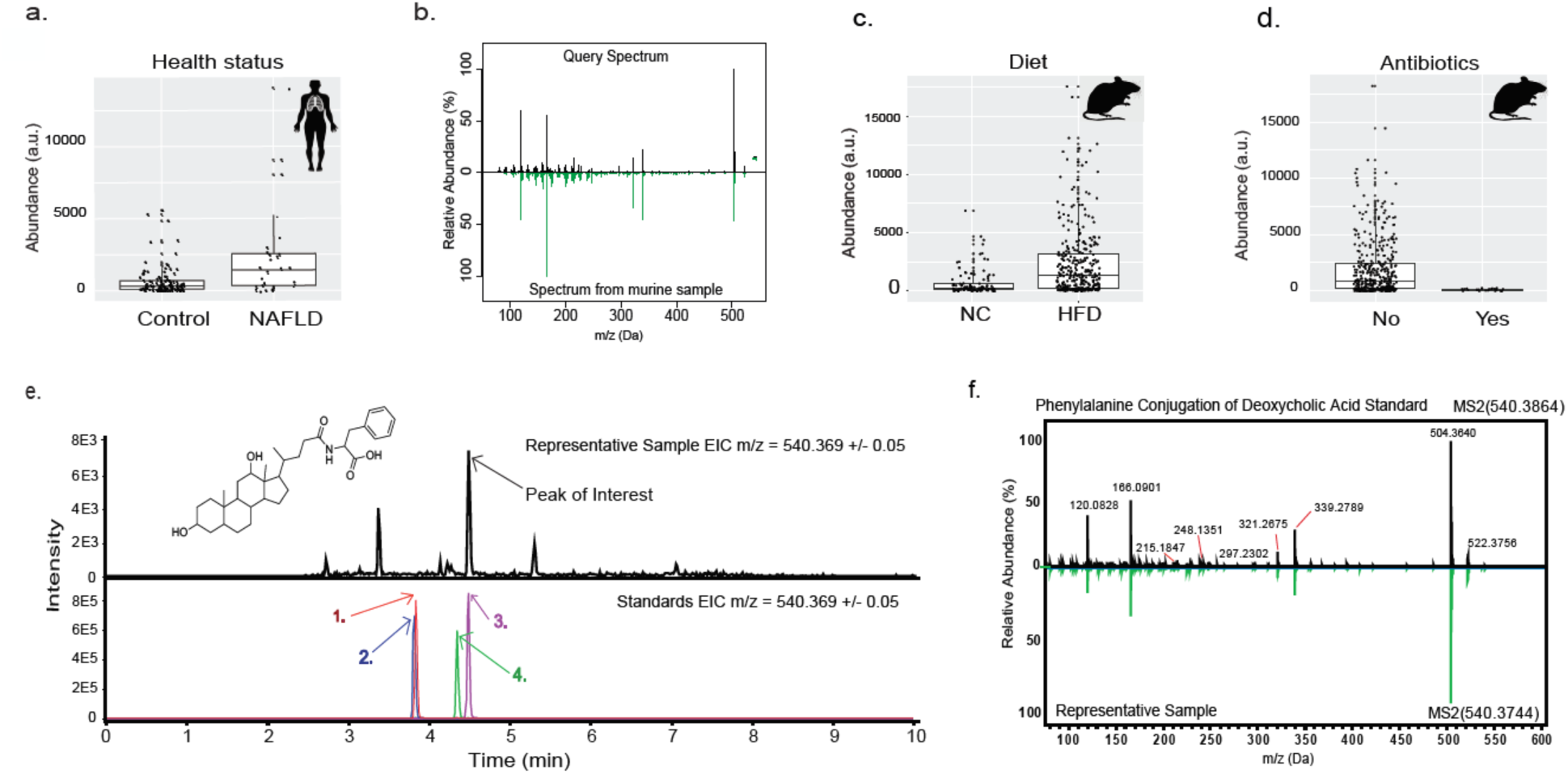
Feature with *m/z* 540.3677 associated with a nonalcoholic fatty liver disease (NALFD). a. Box and whisker plot for the human data, p=9.91E-07 (T-test); b. Mirror plot of the MS/MS spectrum from the NALFD cohort used as MASST query compared to a spectrum found in murine cirrhosis study; c. The levels of this molecular feature in fecal samples from mice fed a high fat diet (HFD) vs normal chow (NC), p=2.7E-08; d. The levels of this molecular feature when mice on a HFD are treated with antibiotics p=7.2E-37; e. Extracted ion chromatograms (EICs) of the feature of interest for a representative sample (top panel) and four synthetic standards (bottom panel, phenylalanine amidate conjugate of 1) ursodeoxycholic acid; hyodeoxycholic acid; 3) deoxycholic acid; 4) chenodeoxycholic acid). f. Mirror plot of MS/MS of the feature of interest observed in a representative sample compared to that of the synthetic standard of phenylalanine conjugate of deoxycholic acid.

**2) Can findings about a molecule identified in model organism studies be translated to humans?** One major application expected will be the translation of molecular information from animal models to humans. The first example of such a translational application was shown during the development of MASST.^2^ In a murine model of acute infection with lymphocytic choriomeningitis virus Armstrong (LCMV ARM)^4^, it was observed that the ileum of infected and uninfected mice contained an MS/MS spectrum with a precursor mass *m/z* 496.326, which was significantly reduced in abundance at day 8 post-infection when compared to uninfected controls (Box Figure 2a). There were neither matches to any reference spectrum, nor could any other match to this molecular feature be found in the metabolomics literature. To test whether this molecule is also found in humans, thus supporting the translational potential, a MASST search was performed. MASST revealed that the same MS/MS spectrum was found in another murine dataset and two human datasets; the American Gut Project (2 males and 7 females all of whom were >45 years of age)^5^ and in four samples of children less than 2 years in an infant eating behavior study. Because this molecular feature was also observed in fecal samples in humans, this molecule was prioritized for structural determination. A molecular network suggested this molecule (Yellow circle, Box Figure 2) was related to a recently discovered set of microbially synthesized bile acids (Green diamonds, Box Figure 2).^2^ Molecular networking suggested that serine was conjugated to cholic acid similar to the phenylalanine, leucine/isoleucine and tyrosine conjugates. Precursor mass shifts between the amino acid conjugates of cholic acid support that serine was conjugated (Box Figure 2). Indeed, comparison of our annotation to a synthetic standard of cholylserine showed identical precursor masses (*m/z*), retention time (RT), and MS/MS spectrum which is level one identification according to the 2007 Metabolomics Standards Initiative (Box Figure 2).^3^

MASST_GNPS job link: https://gnps.ucsd.edu/ProteoSAFe/status.jsp?task=bac3d3788e704af59e4a15a5146e4d6b

Molecular networking job https://gnps.ucsd.edu/ProteoSAFe/status.jsp?task=1809afb755794081b128585409100343

**Box Figure 2.**
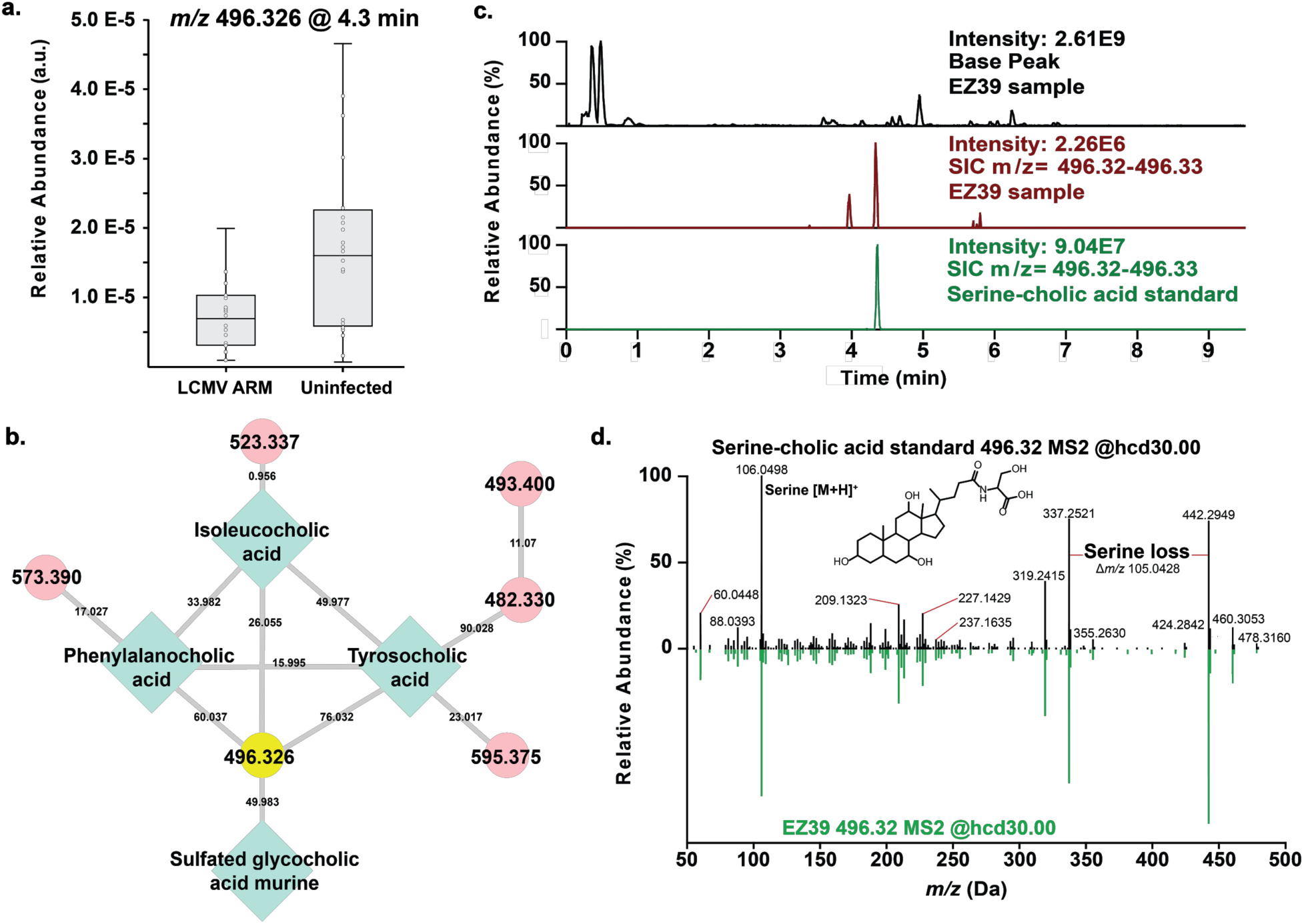
Modulation of a cholylserine bile acid in a mouse model of acute infection with LCMV. a. 10 week-old C57BL/6J mice were intravenously infected with 2×10^6^ pfu LCMV ARM. Untargeted LC-MS/MS and subsequent identification of a molecular feature with an *m/z* at 496.326 was performed in the ileums of infected mice and uninfected controls at day 8 post-infection. The relative intensities were based on the total ion current of that feature. Pooled data from 2 independent experiments are shown. **p<0.01 by Mann-Whitney test. b. The molecular family within the molecular network for this MS/MS spectrum is shown in the aforementioned mice. Green diamonds are nodes with spectral annotations to the reference library, red circleas do not have annotations. Yellow curcle is the node of interests. c. The retention time of the synthetic standard for cholylserine vs a representative uninfected ileum sample are shown. d. The MS/MS spectrum of the synthetic standard compared to the to the MS/MS of the sample (green).

**3) Can MASST be used to reveal the presence and distribution of environmental toxins?** Domoic acid became famous through the novel “The Birds” by Daphne du Maurier and a film from Alfred Hitchcock. This neurotoxin caused seagulls to attack humans. In real life, it has been responsible for numerous poisonings of humans and sea animals, has caused several fatalities^6^, has a negative economic impact such as shutting down crabbing in California and is monitored in many coastal areas on the West and Northeast coast of the United States. From a MASST search, spectral matches to domoic acid were found in seven different public datasets, and although more than 1,000 metabolomics datasets were searched, all seven matches were marine related, including culturing experiments of the diatom *Pseudonitzschia*, one of the known domoic acid producers. Spectral matches to domoic acid were found in datasets from the California coast, including data from the Scripps pier and surrounding beaches in San Diego, an area where domoic acid has been observed frequently. In addition to places where it has been observed, matches to domoic acid were also present in five of the LC-MS/MS runs from data from surface seawater collected in Narragansett Bay, Rhode Island, originally deposited in MetaboLights (MTBLS293)^7^ and a dataset from Hawaiian coral reef water. This is surprising because these areas have few reported *Pseudonitzschia* blooms. Thus, the re-purposing of non-targeted scientific surveys of marine systems, that are costly to generate, might offer a strategy to prioritize regions for expanded toxin monitoring.

MASST_GNPS job link https://gnps.ucsd.edu/ProteoSAFe/status.jsp?task=4d3efb880fda4b8bae36dbfe5b5d6e42

**4) In what datasets can we find a published MS/MS spectrum?** A class of potentially very important compounds has been described in the literature: N-acyl amide lipids were predicted to exist via bioinformatics analysis, and subsequent heterologous expression of a gene cluster from human-associated bacteria demonstrated their existence.^8^ Because of their structural similarity to endogenous signaling molecules of eukaryotes, these compounds are capable of interacting with G-protein-coupled receptors (GPCRs), thus providing a potential way for microbiota to manipulate physiology of the host.^8^ This finding brings about tremendous opportunities for development of new drugs and therapies. Previously, we have revealed the presence of these molecules in human stool samples.^5^ Further MASST search with the MS/MS of 3-hydroxyhexadecanoyl glycine and 3-hydroxypentadecanoyl lysine suggests that these molecules have a very wide ecological distribution. These same MS/MS spectra appear in roughly 10% of all public datasets. In addition to human fecal samples, they were also found in data from fecal samples of mice, rats, human and bovine teeth, isolates of various bacteria including multiple *Streptomyces, Amycolatopsis, Pseudomonas, Neisseria, Achromobacter and Bacillus* species, but also, in multiple marine samples including coral reefs, open ocean waters, marine sediments, as well as soils and even human habitat. Other molecules that have such wide distributions are structurally related molecules such as phospholipids. Perhaps the GPCR response to these molecules is an evolutionary result of microbial co-existence, a hypothesis that was formulated on the basis of searching these published MS/MS spectra.

MASST_GNPS job link:

3-hydroxyhexadecanoyl glycine https://gnps.ucsd.edu/ProteoSAFe/status.jsp?task=c853cad1dee04d25a82ed7d0ad1faf61 3-hydroxypentadecanoyl lysine https://gnps.ucsd.edu/ProteoSAFe/status.jsp?task=f0dbb65d92624d0090b2219dd8789ea8

**5) Are specific natural products observed in cultured microbes also observed in non-laboratory settings?** Many of our therapeutics in use today are derived from microbial natural products.^9^ Many natural products, including natural products widely used in the clinic, have not been detected in environmental samples (as opposed to cultures in the lab) and therefore their ecological roles have not been widely established.^10^ Thus, one question that the natural products research and related fields often consider to answer is whether a molecule found in the laboratory might also be present in natural environments, and if so, where? An example of such a natural product belongs to the viscosin family of pseudomonads derived molecules: orfamides.^11,12^ Several biological activities have been reported for this family of microbial cyclic lipopeptides, supporting their potential as biocontrol agents.^11,12^ While they have been isolated from microbial cultures, they have not been observed in a non-laboratory setting. A MASST search with the MS/MS spectrum of *m/z* 1295.84, corresponding to orfamide A, revealed four datasets that contained this molecular ion. These datasets include not only *Pseudomonas* isolate collections,^36^ but also field-collected *Trachymyrmex septentrionalis* fungus gardens collected from the eastern USA. Ant fungus gardens are intricate multi-microbial systems that are both a home and food source for ants and their larvae.^13^-^16^ These results suggest the presence and, perhaps, an as-yet-undetermined role of pseudomonads in natural ant fungus garden ecosystems. To investigate this hypothesis, we analyzed the publicly available *T. septentrionalis* fungus gardens in NCBI and confirmed the widespread existence of pseudomonads in these environments, consistent with previous work in related systems.^13^-^16^ Furthermore, we isolated several pseudomonads from *T. septentrionalis* fungus gardens. In combination, these findings suggest that orfamides and related molecules might also play a previously unrecognized role in ant fungus garden ecosystems.

MASST_GNPS job link: https://gnps.ucsd.edu/ProteoSAFe/status.jsp?task=bf28bda6ccdd4b699f9a5f2a67b93ee6

**6) Where do we find agricultural fungicides in the environment? Is there evidence that people may be in contact with these fungicides?** Fungicides are widely used in agricultural systems to control for devastating losses worth millions of dollars in crops worldwide. Exposure to fungicides by humans may occur through contact with the skin, ingestion or inhalation.^17^-^20^ A MASST search with the MS/MS spectrum of azoxystrobin, a fungicide of the strobilurin family commonly used in agriculture world-wide revealed matches in expected datasets, including two datasets containing a standard to azoxystrobin, a dataset that sampled the surface of fruits, and several food datasets, especially fruits and vegetables. Deeper inspection of the food data-sets revealed that azoxystrobin was found in a mandarin, pressed juices, herbed goat cheese, restaurant grain dish, cucumber (with peel), roma tomato (with peel), duck, orange juice, peanut butter, red sauce, tomato pesto, dried parsley, muffin, mandarin orange flesh, mandarin orange peel, potatoes with herbs, raw green onion, green grapes and raisin. There was no match to any environmental datasets, but a large number of matches were observed from human skin samples in six different datasets. In total 78 LC-MS/MS files from 15 out of 135 volunteers in all six studies revealed the presence of a spectral match to azoxystrobin from sampling sites included hands, faces and feet suggesting that a significant population might be exposed to azoxystrobin.

MASST_GNPS job link: https://gnps.ucsd.edu/ProteoSAFe/status.jsp?task=f3fbe2baf0be4641bf14e8f60a79494d

**7) Are known toxins from food found in/on people?** The mycotoxin roquefortine C is a potent neurotoxin at high concentrations, but is present at low concentrations in foods such as blue cheeses.^21,22^ The LD50 is 169-189 mg/kg by intraperitoneal administration. Roquefortine C levels ranging from 0.05 to 12 mg kg^-1^ have been reported in cheeses. While it has low toxicity in humans due in part to low bioavailability, the routes of exposure to toxicants and their potential sources are of interest for food safety. As a toxicant exemplar we identified roquefortine C in a number of blue cheese samples. Through a MASST search we also obtained MS/MS matches to two NIH natural products standards reference collections that include roquefortine C as a standard, a *Penicillium* culture, as well as additional blue cheese bacterial and fungal isolates, a fungal collection, and ocean microbial cultures and human stool (infants and adults). The sources and routes of exposure can be thus proposed, beginning with detection in microbial culture, in fermented food products, and ultimately in stool from people.

MASST_GNPS job link: https://gnps.ucsd.edu/ProteoSAFe/status.jsp?task=c1da96a2d4a74568a3394d2dadf2ffff

**8) Can we use approximate matches to a natural product to find datasets that may contain analogs?** Staurosporine is a natural indolocarbazole discovered in 1977 from a culture of *Saccharotbrix sp.* obtained from a soil sample collected in Japan.^23^ A decade after that discovery, its strong cytotoxic effect against cancer cells through its potent modulation of tyrosine kinases was demonstrated.^24^ Here, we leverage the spectra of the staurosporine reference spectrum with MASST search on GNPS to assess its occurrence in the public MS/MS datasets, and to discover potential new derivatives. A MASST search was performed using the staurosporine available in the GNPS library (<u>CCMSLIB00000001655</u>). 14 datasets showed the presence of related spectra, mostly associated with microbes. Spectral matches to staurosporine, were observed in samples originating from four Actinobacteria datasets, which was consistent with previous investigations,^25,26^ but also directly in soil and marine sediments. MASST in analog mode revealed several candidate analogs. Interestingly, amongst these potential analogs, only one spectral match corresponded to previously described derivatives (17-OH staurosporine, +15.98 Da), while all the other annotations were undescribed putative staurosporine derivatives (including +CH_2_, +14.00 Da; +NO, +29.98 Da; +CHN_2_O; +32.99 Da). The search for staurosporine derivatives among the public datasets with MASST took less than 15 min, and suggests that there are still yet to be discovered reservoirs of unique staurosporine derivatives.

GNPS_MASST job link: https://gnps.ucsd.edu/ProteoSAFe/status.jsp?task=e5b04be34a2f41cb9b681e472d9a1765

**9) Can MASST be used to track sunscreens in human and environmental samples?** Understanding the impact of humans on earth’s ecosystems is of increasing importance. Sunscreens became part of the formulations of several skin care products and they are used worldwide on a regular basis, to protect the skin from the damaging effects of ultraviolet radiations including sunburn and skin cancer. ^27^ However, when it leaves our skin where does it end up? A MASST search of the MS/MS spectra of two active ingredients of sunscreen - avobenzone and octocrylene – reveals, as expected, their presence in many human skin datasets. avobenzone and octocrylene were detected in skin samples from 19 public datasets, including 15 datasets from the United States, 2 datasets from surfers from Morocco and England, 1 dataset from Japan and 1 dataset from individuals living in Venezuela. Additionally, avobenzone and octocrylene were detected in saliva, teeth and stool. While skin photoprotection is crucial for human health, many questions have been raised regarding the accumulation of sunscreens in the environment. ^27^-^30^ To look for sunscreen in environmental samples, MASST found matches to the indoor environment or personal objects such as offices, houses, cars, bikes, phones, wallets, keys, mattresses, plants, meat for human consumption, corals, and even in coral reef in remote areas such as Moorea.^31^-^33^ While there have been no identifiable toxic effects on humans ^34^, these results show that human-made chemicals may be widely distributed without truly understanding the potential long-term impact.

Avobenzone *m/z* 311.165 MASST_GNPS job link: https://gnps.ucsd.edu/ProteoSAFe/status.jsp?task=56a764c64f524a30b5796f5d9124b832

Octocrylene *m/z* 362.211 MASST_GNPS job link: https://gnps.ucsd.edu/ProteoSAFe/status.jsp?task=84346268fdca4144bf71d9cdb42fef02

**10) Can we find evidence of opioids exposure in public data?** Opioids are used not only as psychotropic drugs but are also prescribed as analgesics for the relief of severe pain. In the US, the use of these substances often begins with a prescription for pain relief.^35,36^ Methadone is a synthetic opioid used to treat drug addiction and is also used in palliative care for patients with cancer, HIV and postoperative related pain.^36^ By searching the MS/MS of methadone using MASST, we found a matching MS/MS spectrum in five datasets. Methadone was detected in stool samples of two subjects from the American gut project dataset, both are males, one is middle adulthood and the other is early adulthood. It was found in 5 patients affected by inflammatory bowel disease (IBD). Those patients suffer from chronic abdominal or musculoskeletal pain and opioids (including methadone) are commonly prescribed as pain relief.^35^ 12 of the samples of the ∼1000 teeth contained a spectral match to methadone. This observation is supported by the fact that dentists in the US, commonly prescribe opioids as analgesics, particularly for surgical tooth extraction.^36,37^ However, it is worth highlighting that two of the teeth samples also matched reference spectra of cocaine; the co-occurrence suggests the rationale for methadone could stem from recreational use or relapse of treatment. Finally, in addition to human samples, we found methadone in one sediment and four water samples belonging to a dataset called Earth Microbiome Project (EMP)_500_Metabolomics, a study on environmental samples collected throughout the US.^38, 39^ This suggests that methadone (and likely other drugs) are present in our environment.

Methadone MASST GNPS job link: https://gnps.ucsd.edu/ProteoSAFe/status.jsp?task=afa2d37fddb44af2ab016c4c7493b3fc

Cocaine MASST GNPS job link: https://gnps.ucsd.edu/ProteoSAFe/status.jsp?task=c40a73f588b24975b541788d7f086510

